# Domain-adaptation deep learning models do not outperform simple baseline models in single-cell anti-cancer drug sensitivity prediction

**DOI:** 10.64898/2026.02.24.707713

**Authors:** Michael Bohl, Marina Esteban-Medina, Niko Beerenwinkel, Kerstin Lenhof

## Abstract

Tumor drug response is profoundly shaped by cellular heterogeneity, making single-cell resolution essential for precision oncology. While drug-response labels are abundant for cell lines at bulk resolution, translating these predictive models to the single-cell level requires effective domain adaptation strategies. Motivated by advances in computer vision, recent deep-learning domain adaptation methods promise to transfer knowledge from bulk (source) to single-cell (target) data without the need for target labels. However, their true translational utility remains unclear due to a lack of rigorous evaluation against non-adaptive baselines across diverse biological and technical contexts. Here, we present a comprehensive benchmark comparing four representative domain adaptation methods against two simple gradient boosting baseline methods. Through systematic evaluation across 19 single-cell datasets and 10 drugs, we show that none of the complex adaptation methods outperforms the simpler baselines. By analyzing the drivers of model performance, we find that target-informed hyperparameter tuning and sparse label supervision are the principal sources of prediction gain. Our study reveals that current approaches fail to bridge the bulk-to-single-cell conceptual shift and provides a unified codebase and comprehensive data collection to facilitate robust model comparisons. By enabling transparent evaluation and robust benchmarking against simple models, this resource aims to accelerate future developments in translational pharmacogenomics.

## 1 Introduction

A primary goal of personalized medicine is to tailor treatments to patients based on their phenotypic and genotypic characteristics. In personalized oncology, the optimization of drug therapy based on molecular tumor data to improve patients’ response is of particular interest. To achieve this aim, various machine learning (ML) methods for predicting the effectiveness of anti-cancer drug treatments have been developed [1–4]. Most of these methods are trained on data from model systems, typically cell lines, since such model systems can be relatively easily exposed to the existing multitude of approved and experimental anti-cancer drugs [1, 5]. Here, gene expression profiles obtained from high-throughput RNA sequencing or microarray experiments constitute the feature space [6–10]. Importantly, these gene expression values are bulk measurements; that is, they reflect averages across a population of cells. The response variable is defined through a dose–response curve and usually summarized by IC50 [11] or Cmax viability [12] values.

A central hypothesis driving method development is that models derived from these scalable cell line experiments can be translated to more complex biological systems, such as patient-derived xenografts, ultimately guiding treatment selection for human tumors. Recent efforts have focused on transferring the learning from bulk cell line-based models (source domain) to predict responses in increasingly complex model systems, up to patient samples at the single-cell resolution (target domain) [5]. This translation into the single-cell domain is key for precision oncology, as different subpopulations of tumor cells, in general, respond differently to treatment [13–15].

However, this transition is hindered by a substantial domain shift encompassing profound biological differences (homogeneous cell lines vs. complex tissues), measurement platform discrepancies (bulk RNA-seq vs. scRNA-seq), and severe annotation gaps (fully labeled source vs. sparsely labeled target) [7, 9]. To bridge this gap, recent methods have adopted domain adaptation deep learning strategies inspired by computer vision [16–21].

Current frameworks adopt two main strategies: Unsupervised Domain Adaptation (UDA), which learns domain-invariant representations using only labeled source and unlabeled target data, and Semi-Supervised Domain Adaptation (SSDA), which incorporates a small set of labeled target samples to guide alignment. To evaluate the current landscape, we selected four state-of-the-art methods that exemplify the distinct algorithmic paradigms. SCAD is based on an adversarial approach, which aligns bulk and single-cell representations via a generative adversarial objective to learn domain-invariant features for drug response prediction [22]. scDEAL represents a metric-based method that minimizes maximum mean discrepancy (MMD) between bulk and single-cell latent spaces to transfer drug sensitivity knowledge while preserving cellular heterogeneity [23]. Representing the emerging use of foundation models, scATD leverages pretrained scFoundation embeddings and knowledge distillation to guide MMD-based domain alignment [24, 25]. Finally, SSDA4Drug employs an SSDA strategy that alternates between entropy maximization and minimization to exploit a few labeled target cells [26].

Despite the proliferation of such methods, their true translational utility remains unclear. Existing studies often differ in evaluation metrics, data splits and supervision protocols. In addition, they often provide limited or inconsistent comparisons against simple baselines, leaving the question of whether increasingly complex architectures deliver genuine advantages beyond what simpler models can achieve. Moreover, none of the studies addresses inherent data set issues, e.g., that many single-cell datasets label untreated cells as sensitive and treated cells as resistant, thereby misclassifying pre-existing resistant subclones and exaggerating class separation by incorrectly attributing treatment-induced transcriptional changes to resistance.

To address these limitations, we carried out a comprehensive benchmarking study that systematically evaluates bulk-to-single-cell drug-sensitivity prediction deep learning methods and tests whether they achieve meaningful domain adaptation by comparing them against simpler baselines. We curated a comprehensive set of 19 singlecell drug-response datasets across 10 drugs to compare the four representative methods SCAD [22], scDEAL [23], and scATD [24], alongside SSDA4Drug [26]. In addition, we included two gradient boosting methods as baselines: a CatBoost [27] model trained only on bulk data (unsupervised regime) and a CatBoost-few-shot model trained on a few labeled single cells per class (few-shot regime). Since these baselines rely solely on standard supervised learning and do not attempt explicit domain adaptation, they establish the transfer performance achievable without distribution alignment.

Our work reveals that current deep learning domain adaptation methods do not yet achieve meaningful bulk-to-single-cell translation. Unsupervised methods (SCAD, scDEAL, and scATD) perform close to random when strictly tuned on source data. Our analyses suggest that their previously reported performance likely relies on target-informed hyperparameter tuning. Furthermore, a standard CatBoost model matches these complex architectures when tuned similarly. Most notably, a simple few-shot CatBoost baseline, which uses few target labels but lacks any domain-alignment strategy, matches or outperforms all evaluated state-of-the-art methods while providing superior efficiency and interpretability.

The proposed benchmark provides a transparent basis for comparing complex approaches with simpler baselines and determining when performance reflects genuine cross-domain transfer versus target-informed optimization. To support reproducible progress in bulk-to-single-cell drug-response prediction, we have made the benchmark and evaluation protocols available at https://github.com/cbg-ethz/SC-Bulk-Domain-Adaptation.

## 2 Results

To determine whether deep learning–based domain adaptation methods offer advantages over simpler baseline models, we compiled a comprehensive benchmarking collection of 19 single-cell datasets across 10 drugs exceeding, to the best of our knowledge, the scale of previous comparisons. We employed harmonized preprocessing, stratified splits, and consistent assessment protocols (for details, see Methods 4.4) for our comparative assessment of methods predicting single-cell drug response from bulk-trained models. We re-implemented the UDA methods SCAD, scDEAL, and scATD, as well as SSDA4Drug within a unified training environment, exactly matching the architectures, loss functions, and optimizers of their original implementations. For each drug, we trained models on labeled source (bulk RNA-seq and microarray) data and unlabeled target (single-cell RNA-seq) data and evaluated them on previously unseen samples from the source and target samples in a binary classification task (sensitive vs. resistant; see Fig. 1 and Methods). In addition, we assessed model performance on held-out target datasets, i.e., single-cell data from the same drug but generated under different experimental conditions, and, crucially, entirely excluded from training. This setup allowed us to quantify the ability of each method to generalize beyond the particular single-cell target data set available during training.

**Fig. 1:**
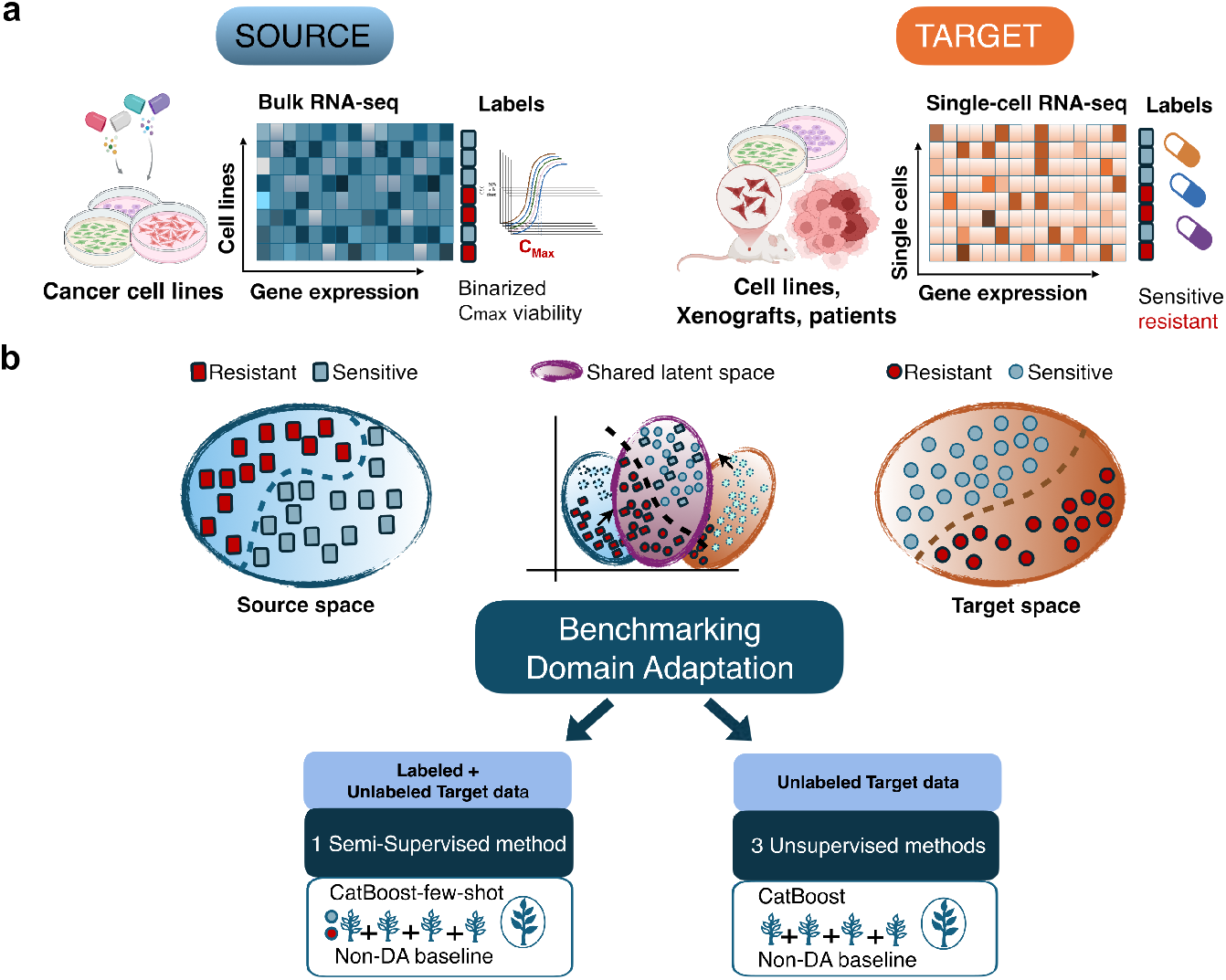
Overview of the bulk-to-single-cell domain adaptation benchmarking study for drug sensitivity prediction. (a) Source and target domains: cell-line bulk RNA-seq profiles with dose-response viability curves (normalized by *C*_max_ viability) are converted into binary sensitivity labels, while the target domain contains single-cell RNA-seq profiles from cell lines, xenografts, or patient samples with proxy-derived binary labels. (b) Methods and evaluation: domain adaptation methods align source (squares, top left) and target (circles, top right) data in a shared latent space (top middle) and are compared with non-DA CatBoost baselines in semi-supervised (bottom left) and unsupervised (bottom right) settings.

We first examine the efficacy of domain adaptation strategies, beginning with the impact of target-informed tuning on UDA models and compare them to non domain-adaptive baselines. We subsequently assess SSDA architecture against a fewshot baseline, investigate biases introduced by treatment-proxy labeling, and test the transferability of bulk source optimization to single-cell targets. For these analyses, “target performance” refers strictly to unseen samples from the target dataset used during training, prior to our final evaluation of model generalization on fully independent, held-out single-cell datasets. We report both AUROC and the Matthews correlation coefficient (MCC) to ensure a robust evaluation of real-world performance across concrete decision thresholds.

### 2.1 Unsupervised gains rely on target-informed model tuning

In unsupervised domain adaptation, hyperparameters need to be chosen without using labeled target data, i.e., target labels are reserved exclusively for evaluation. To disentangle true domain-invariant learning from target-specific tuning, we evaluated both: a source only hyperparameter tuning and an intentionally optimistic target-informed setting in which hyperparameters were selected based on target-test performance. We refer to this latter setup as target-informed hyperparameter tuning. We find that in the target-informed hyperparameter tuning, AUROC values are comparable to those reported in the original publications (Fig. 2b). In contrast, when the same models are tuned solely on source domain validation data (Fig. 2c), the target domain AUROC values decline substantially, approaching random performance (the mean AUROC hovered around 0.5, and the mean MCC approached zero across all 19 datasets). This finding indicates that feedback from labeled target samples might be the main performance driver of UDA models rather than domain-invariant representations.

**Fig. 2:**
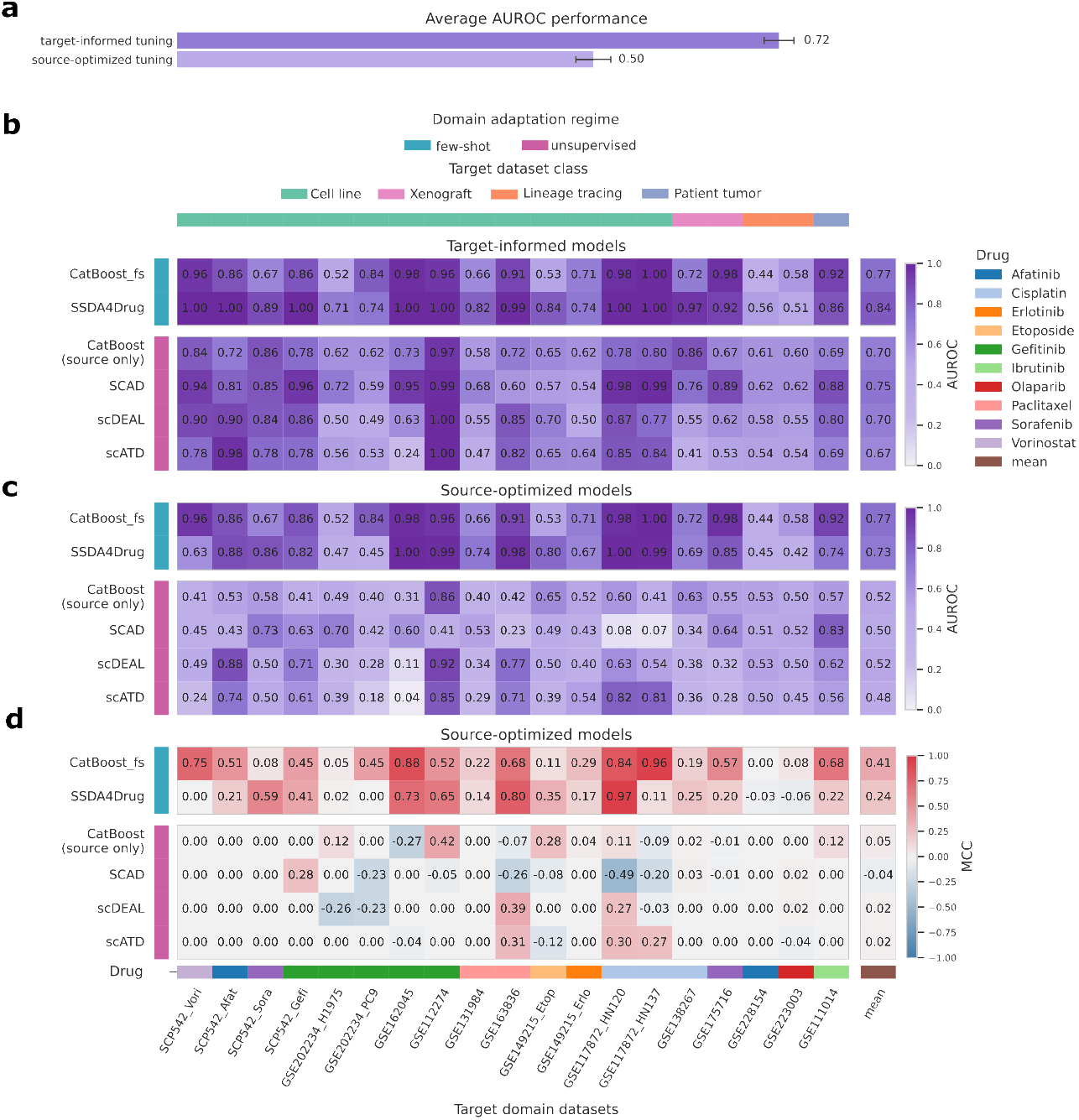
Target domain performance with hyperparameters tuned on either source or target. (a) Mean AUROC of UDA models across all drugs, with hyperpa-rameters tuned either using the source domain or the target domain. The heatmaps show (b) AUROC for each model–dataset pair with target-optimized hyperparameter selection; (c) AUROC for each model–dataset pair with source-optimized hyper-parameter selection; (d) MCC evaluated at a single decision threshold chosen on source-validation data. Columns correspond to single-cell target datasets, except for the final column (mean), which reports the average performance per drug, computed by first averaging across datasets belonging to the same drug and then averaging across drugs. Rows are evaluated prediction methods. The top annotation bar encodes target sample type and the bottom annotation bar encodes the drugs tested in the different target data sets. The row annotation bar indicates the domain-adaptation regime (few-shot vs. unsupervised). Higher (darker) is better for both performance metrics; random predictions correspond to MCC = 0 and AUROC = 0.5.

Moreover, we find that performance varies across drug–dataset pairs: isolated high performance on individual datasets did not replicate across other datasets for the same drug, suggesting potential dataset-specific effects rather than systematic drugresponse transfer. In most cases, the AUROC of scDEAL and scATD showed a strong correlation (Spearman *ρ* = 0.86). In contrast, SCAD performance outliers occurred across largely distinct datasets, indicating that these methods learn dataset-specific patterns rather than generalizable drug-response features (Fig. 2bc). Overall, despite explicitly optimizing for domain-invariant features, no UDA model’s average AUROC or MCC surpassed the source-only CatBoost baseline (Fig. 2cd, last column).

### 2.2 Simple few-shot baselines match semi-supervised domain adaptation

We further compared SSDA4Drug to a simple few-shot baseline: a shallow Cat-Boost classifier (30 trees, depth 3) using six labeled target cells (three per class, the same number of target labels provided to SSDA4Drug) in addition to source domain data. Despite additionally leveraging unlabeled target samples and source supervision through its semi-supervised objective, SSDA4Drug did not outperform this baseline on average (mean AUROC and MCC are 0.73 and 0.24 for SSDA4Drug and 0.77 and 0.41 for few-shot CatBoost, see Fig. 2cd). Moreover, there was no consistent pattern regarding characteristics of the datasets for which SSDA4Drug or the few-shot baseline performed better: there was no clear association with specific drugs (Kruskal-Wallis H-test; p = 0.893), label imbalance (Spearman correlaion; p = 0.718), or the number of genes shared between source and target datasets (Spearman correlaion; p = 0.597). Supplementary Table 5 shows more non-significant associations between dataset attributes and predictive performance.

### 2.3 Labeling sensitivity by treatment status inflates model performance

Most single-cell datasets used in prior UDA/SSDA benchmarks utilize treatment status as a proxy for sensitivity labels (untreated = sensitive; treated = resistant) or assign labels by selecting extreme phenotypes based on marker expression, where a marker (e.g., EpiSen) denotes a predefined gene-expression program and only cells in the top and bottom 10% of marker activity are labeled as sensitive or resistant, with all intermediate cells discarded. The only two exceptions, where neither of these two labeling schemes was used, are the two primary tumor datasets (GSE111014 and GSE169246) included in the scATD evaluation [24]. The above-mentioned treatment-based label definition often introduces some form of shortcut learning, where the model learns the experimental condition in the single-cell expression space rather than the underlying biology of drug sensitivity (see Figure 3). In contrast, lineage-tracing experiments, which utilize clonal barcoding to identify intrinsically resistant subpopulations prior to treatment, offer a more clinically grounded framework for target prediction. For these lineage-tracing datasets (orange annotation in Fig. 2), all models exhibited substantially degraded performance. This trend was particularly evident for the drug Afatinib, where model performance diverged between a lineage-tracing dataset (GSE228154, Fig. 3a) and a dataset with extreme phenotype labeling (SCC47, Fig. 3b). Using target-informed hyperparameters did not improve performance for the lineage-tracing dataset (AUROC ≤0.61); it remained considerably below that observed on the extreme-phenotype dataset (AUROC ≥0.78; Fig. 2). Another illustrative example of such separability are the GSE117872 datasets HN120 and HN137, where resistant and sensitive cells form almost perfectly distinct clusters in expression space (Fig. 3d). In this setting, the near-complete class separation means that many decision boundaries, will separate the two classes, even if they do not reflect meaningful drug-response patterns. This structure translates directly into model behavior: scATD achieves AUROC values above 0.8, whereas SCAD performs almost perfectly inversely (AUROC ≤0.1), indicating that extreme separability can yield either seemingly excellent or poor performance, depending on whether the decision boundary assigns the two clusters correctly or in reverse, rather than true predictive accuracy. When a few target labels are available, SSDA4Drug and even the shallow few-shot CatBoost readily achieve AUROC ≥ 0.98 on these datasets (Fig. 2b).

**Fig. 3:**
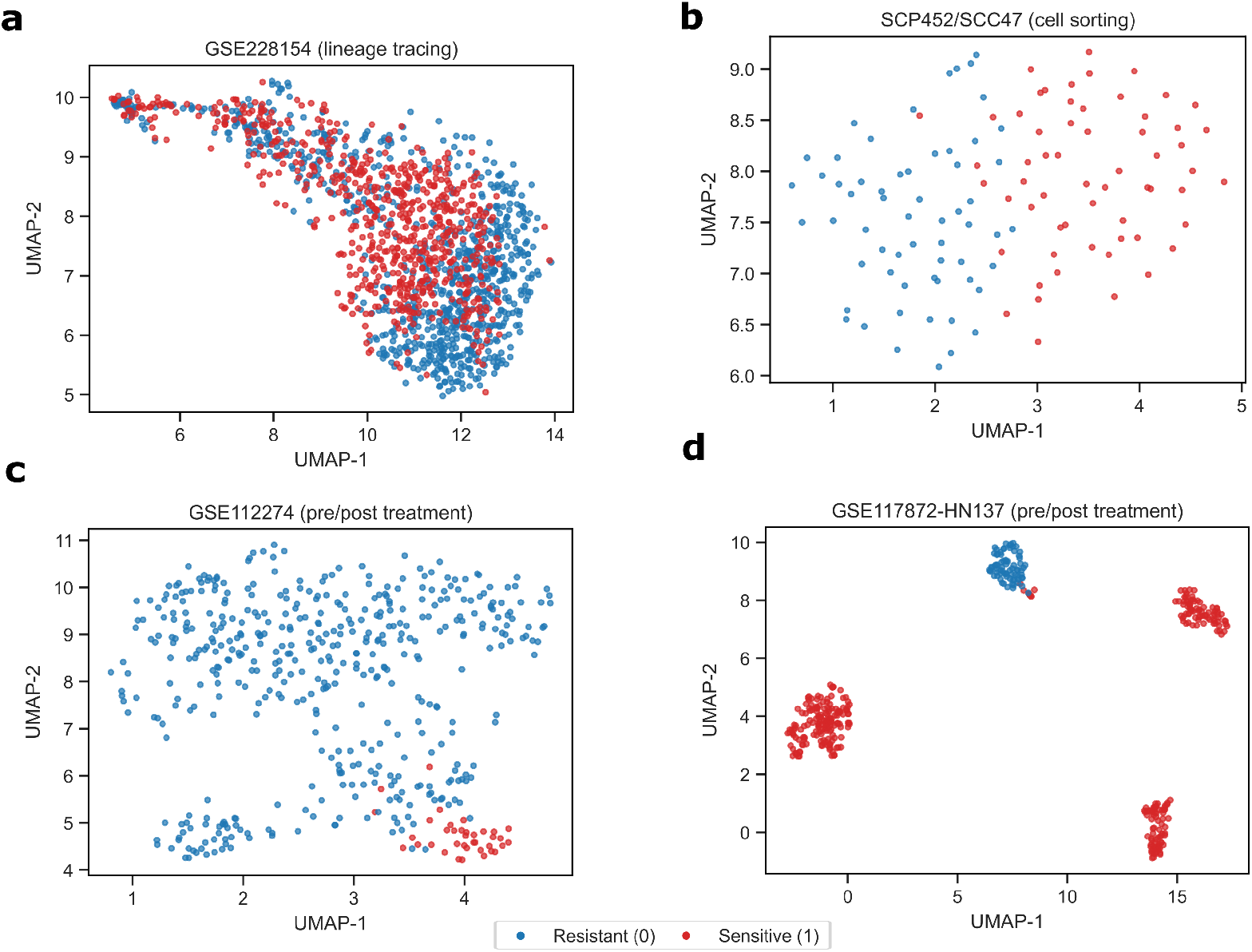
Example UMAPs of single-cell Single-cell datasets with and without inflated separability. (a) Lineage tracing dataset, labeled with Afatinib resistance. (b) SCC47 dataset, including only cells with the most extreme EpiSen marker levels, used in the SCAD and SSDA4Drug publications. EpiSen marker status was used as a proxy for Afatinib resistance. (c,d) Datasets with labels assigned based on treatment status used in both scDEAL and scATD where Cisplatin resistance labels were assigned based on treatment status.

### 2.4 Impact of source domain optimization on target predictive performance

To assess whether strong performance on the source domain translates to improved target domain performance, we conducted Bayesian hyperparameter optimization as implemented in Optuna [28], with 100 trials per model–dataset pair (10,500 runs in total), optimizing source-validation MCC while recording source-test and target-test metrics. Models were optimized using the source-validation MCC, while performance on the source-test and target-test sets was recorded (see Methods). Source domain optimization failed to identify models that generalize to the single-cell target domain, with no consistent relationship between source and target performance across architectures (Figure 4ab). As expected, hyperparameters that improved source validation AUROC also increased source-test AUROC (Fig. 4c) and MCC (Supplementary Fig. 9a). However, this improvement did not carry over to the single-cell target domain (Fig. 4d and Supplementary Fig.9b). All domain adaptation methods outperformed the CatBoost few-shot baseline on source data (trained and evaluated), but the advantage vanished entirely in the target domain. SCAD, the simplest UDA architecture, showed the largest boost in source domain AUROC after tuning; yet, this gain did not translate to the target domain, underscoring that better source domain optimization does not imply effective domain alignment. The estimates in Fig. 4a show that across all domain-adaptation methods, there was no positive association between source-validation and target-test AUROC, demonstrating that source-only criteria could not reliably select models that generalize to the target domain in our experiments.

**Fig. 4:**
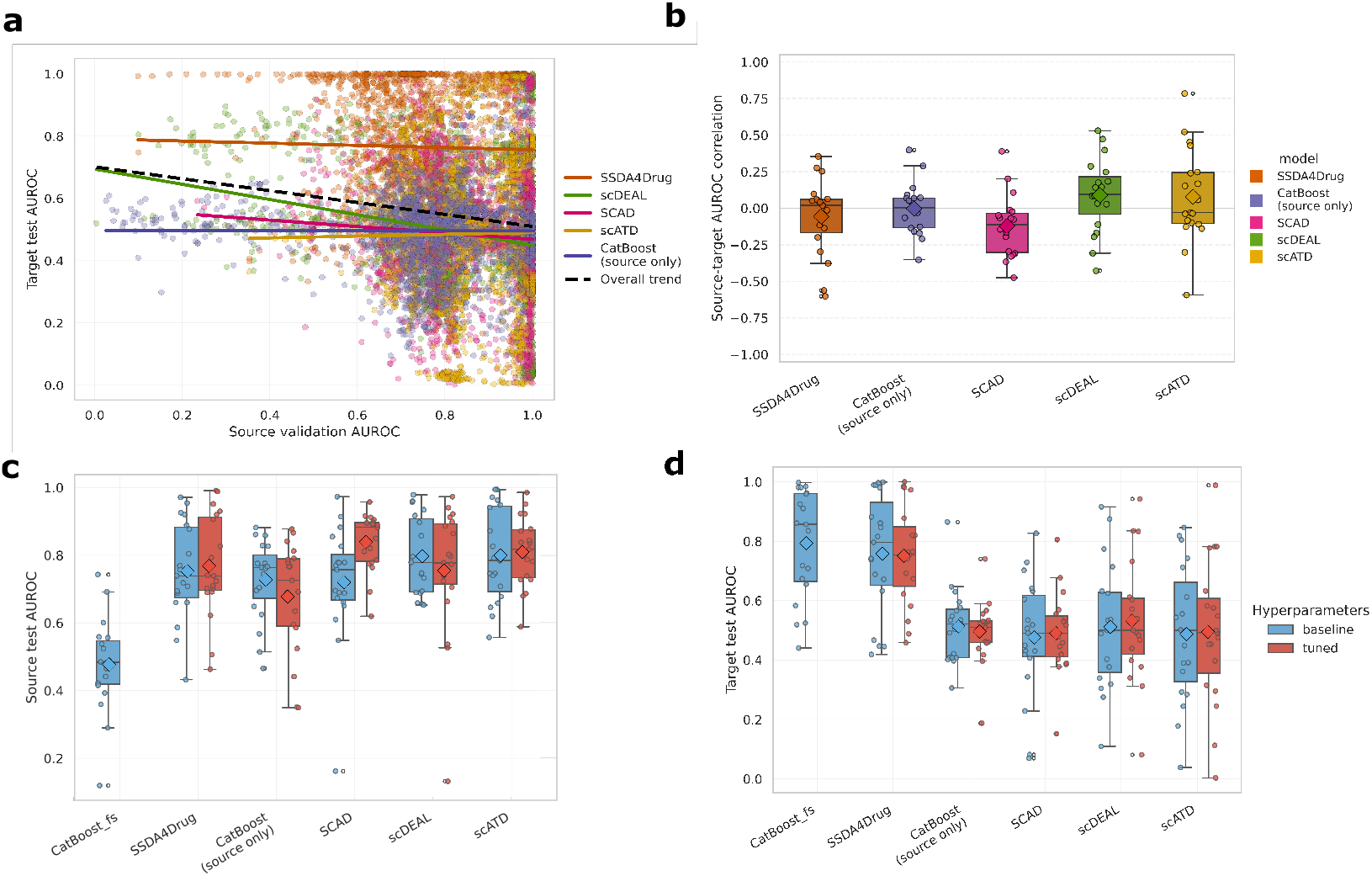
Source-based hyperparameter tuning fails to improve target performance. (a) The relationship between target and source test AUROC scores is illustrated across 1,900 hyperparameter combinations and 19 datasets for each model.(b) Boxplots displaying the distribution of Spearman correlation coefficients between source and target AUROC for individual datasets, with diamonds representing the mean correlation. The distribution of performance across all datasets is further detailed for each model using (c) source-test (bulk) AUROC scores and (d) target-test (single-cell) AUROCs, where diamonds denote the mean performance in each category.

### 2.5 Models fail to generalize across independent datasets of the same drug

A robust transfer model should yield consistent predictions when applied to new, independent datasets of the same drug. We tested this cross-dataset validity using six drugs that appeared in multiple target datasets (Table 4). Despite predicting sensitivities to the same treatments, performance was highly unstable. Models that perform well on one dataset frequently performed no better or worse than random guessing on another when comparing AUROC/MCC scores (Fig. 5). This inability to generalize was most evident for Gefitinib: We utilized three independent studies (GSE112274, GSE162045, and GSE202234) that treated the same “PC9” cell line but differed in laboratory protocols and treatment conditions. This allowed us to evaluate six pairwise combinations of a target dataset: training on one dataset (used for training/validation) and testing on a strictly independent one used solely for testing. As shown in Fig. 5a, models failed to overcome technical variations in laboratory protocols. Instead of learning the biological response to Gefitinib, the models overfit to dataset-specific artifacts. Consequently, the mean performance across all combinations remained at chance levels (Mean MCC = 0, Fig. 5c). Even the highest-scoring methods, SCAD (mean MCC: 0.045) and the source-only CatBoost (mean AUROC: 0.548), demonstrated poor generalizability.

**Fig. 5:**
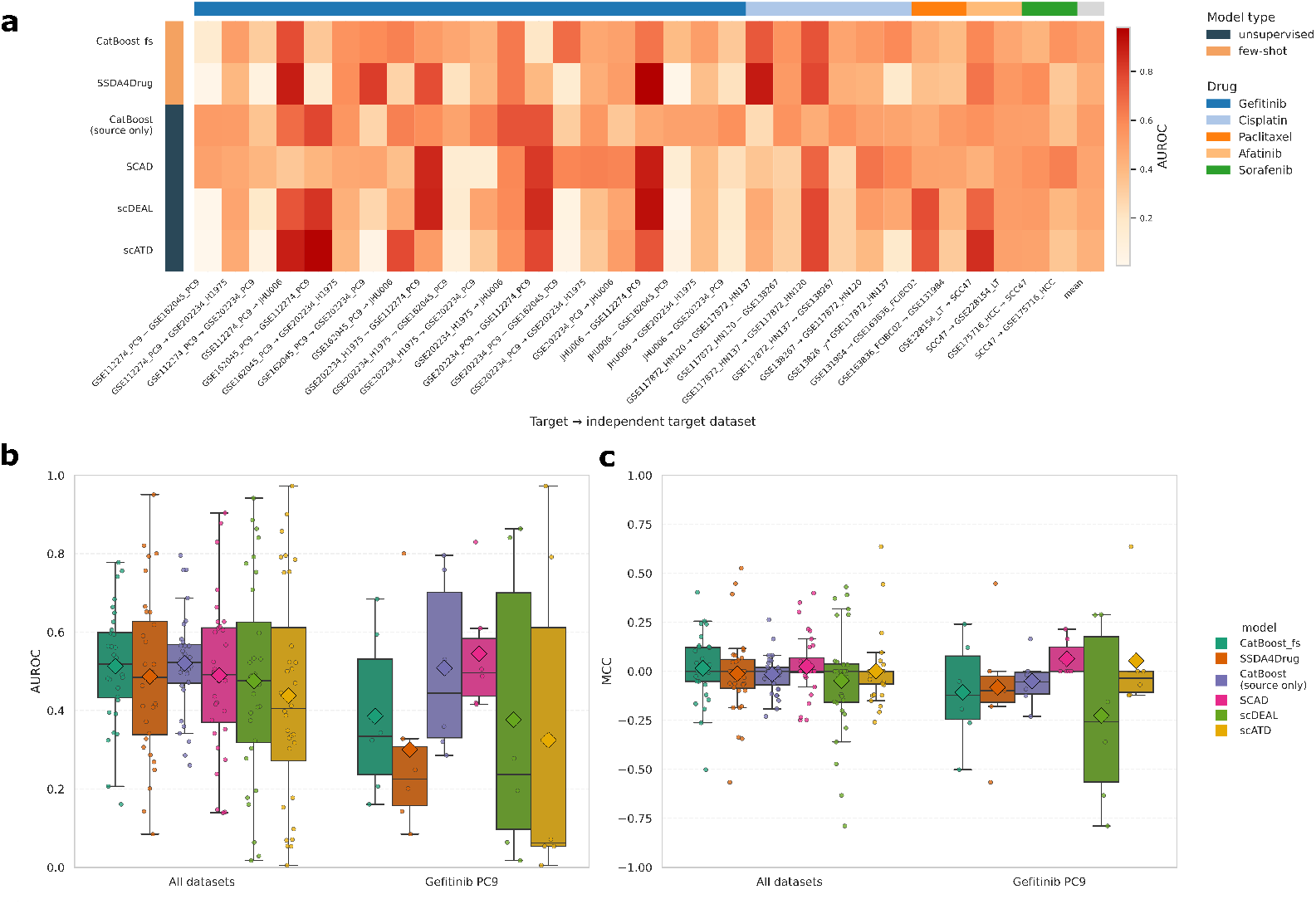
Cross-dataset generalization performance. (a) AUROC scores for all possible target/independent target pairs in our data. Columns are target/independent target pairs, except the last column (mean), which is the mean score across drugs and target datasets; rows are evaluated methods. The annotation bar on the top indicates the drug treatment and the vertical annotation bar on the left indicates the model type. (b) AUROC and (c) MCC comparing the performance of models trained and evaluated on held-out single-cell datasets with response labels for the same drugs. Left: Performance on all 32 train/held-out datasets with heterogeneous cell biology. Right: Performance on train/held-out datasets using the same cell line (PC9) and sensitivity labels for the same drug (Gefitinib). Diamonds indicate the mean.

## 3 Discussion

In this study, we present a unified benchmark to systematically evaluate domain-adaptation strategies for predicting drug responses in individual cancer cells from single-cell gene expression profiles, using models trained on bulk transcriptomic data. We compare state-of-the-art neural network approaches with gradient-boosted baselines in a harmonized framework that standardizes preprocessing, data splits, and hyperparameter tuning. This setup allows us to disentangle transfer-learning gains from improvements driven by other evaluation artifacts.

Our results indicate that existing UDA methods do not currently deliver robust bulk-to-single-cell transfer under realistic conditions. We observed that previously reported performances, as described in the original publications, seemed to rely on hyperparameter selection guided by target domain performance. In a strictly unsupervised setting, where model selection relies solely on the source domain (bulk data) validation split results, UDA methods frequently degrade to random-chance performance when predicting in the target domain (single-cell data). Our evaluations, using the class-imbalanced-aware MCC metric, revealed that modest gains in the AUROC ranking metric (0.6–0.7) often fail to translate into usable decision boundaries, yielding near-zero MCCs (random prediction).

Furthermore, while Semi-Supervised Domain Adaptation (SSDA) methods appear to be superior, our results suggest that this performance is a consequence of partial supervision rather than domain-invariant representations of source and target data. To disentangle these factors, we compared SSDA against a few-shot CatBoost baseline as a non-adaptive control. The few-shot CatBoost operates in the unaligned feature space and is exposed to the same number of target labels as the SSDA model. The fact that the non-adaptive baseline matched the performance of the complex SSDA architecture demonstrates that the adaptation components contribute little to the final accuracy. The improvements relative to UDA are attributable solely to the explicit supervision from target labels, indicating that simpler models can leverage equal effectiveness while remaining more interpretable and computationally efficient.

A recurring pattern in our experiments is that explicit domain alignment often harms performance relative to non-adaptive baselines — a phenomenon known as negative transfer. This suggests that the fundamental assumption of standard domain adaptation, namely that the domain shift is primarily a covariate shift, is violated in this biological context. Most domain-adaptation algorithms, originally inspired by computer vision, assume that while the marginal distribution of features changes, the conditional relationship between features and labels remains stable. In molecular biology, however, the transition from bulk to single-cell data induces a profound concept shift. This shift arises because a bulk expression vector represents a population average (*E*[*X*]), smoothing over stochastic noise and cell-cycle variability, whereas a single-cell vector captures a noisy, high-dimensional snapshot of a specific cellular state (*X*_*i*_). Consequently, the fundamental biological rules mapping expression to drug sensitivity labels differ between the two modalities. Furthermore, this alignment problem is exacerbated by a structural asymmetry between the domains. The source domain (bulk samples, e.g., the GDSC dataset) exhibits high global heterogeneity at low resolution, comprising expression vectors from over 1,000 distinct cancer cell lines that represent diverse tissues and biological contexts. Conversely, the target domain consists of high-resolution single-cell data that typically captures a restricted biological scope; even a heterogeneous patient tumor generally contains a limited number of distinct clonal populations. This structural asymmetry introduces a specific challenge during domain adaptation. Without proper calibration, the alignment process may erroneously force the narrower single-cell target distribution to match the broader variance of the source latent space. Such an over-alignment imposes artificial heterogeneity onto the singlecell representations, distorting their underlying biological structure and degrading the performance of downstream drug-response predictions.

When domain adaptation methods strive for domain invariance, they effectively force the detailed intra-sample heterogeneity of the single-cell target to map onto the broad inter-sample diversity of the bulk source. This process risks collapsing the fine-grained distribution of individual cellular states onto a manifold defined by distinct biological backgrounds, aligning the specific signal of the single-cell data to match the global variance of the bulk population. Attempting to align these distinct modalities without explicitly modeling the generative differences or the hierarchical relationship between a population and its constituent cells likely results in the loss of biological signal and the observed negative transfer effect.

Our findings should be interpreted within the context of several limitations that indicate avenues for future refinement. First, to maintain computational feasibility across a large-scale benchmark comprising over 1,900 hyperparameter combinations, we relied on a single train-test split per dataset rather than full cross-validation. While we mitigated this constraint by evaluating performance across multiple independent datasets, future work should include cross-validation to better estimate variance. Second, our few-shot experiments utilized a single random seed for selecting labeled examples; given that performance can be sensitive to the specific cells selected for supervision—as observed in SSDA4Drug—multi-seed averaging would provide a more robust estimate of stability in low-data regimes. Third, we applied strict inclusion criteria, selecting only drug-datasets for which *C*_*max*_ viability data was available to ensure high-quality ground truth labels, a necessity that restricted our coverage by excluding many otherwise potentially usable datasets with broader drug coverage. Finally, our benchmark reflects the inherent biases of public data repositories, resulting in an over-representation of specific cancer types, such as lung and breast adenocarcinoma, and well-studied compounds. Thus, the generalizability of these findings to rare cancers or novel therapeutic agents remains an open question for future investigation.

Despite these challenges, the establishment of a rigorous benchmarking framework is a critical step forward. Our proposed pipeline provides a modular, metric-rich environment designed for transparent comparison, enabling standardized and reproducible model comparisons. By open-sourcing these tools, we aim to facilitate the integration of diverse datasets and the rapid prototyping of new methods.

Ultimately, this study suggests rethinking single-cell pharmacogenomics. Future progress will likely not stem from merely increasing the complexity of model architectures, but rather from considering the fundamental principles of domain adaptation itself. The next generation of models must move beyond rigid statistical alignment to capture the nuanced biological relationship between broad tissue-level patterns and granular cellular states. By addressing this translational challenge, we can move closer to the goal of precision oncology, fostering models that are better equipped to predict therapeutic responses with biological fidelity.

## 4 Methods

We benchmark four domain adaptation methods that predict single-cell drug responses (unlabeled target domain) from labeled bulk cell line data (labeled source domain). Both domains share gene-expression features but differ in their biological and technical distributions, reflecting the shift from bulk to single-cell. The domain adaptation methods we compare can be categorized into two types of learning settings: UDA, which relies only on labels from the source domain, and SSDA, which uses a few labeled target samples to guide transfer. Each model jointly optimizes a prediction loss on source (and optionally few-shot target) data with an alignment loss that reduces feature-space divergence between domains, aiming to learn domain-invariant discriminative representations that generalize from bulk to individual cells. In the following, we briefly explain how to mathematically formulate this problem and how the four domain adaptation methods address it.

### 4.1 Problem statement

Let 𝒳^*s*^, 𝒳 ^*t*^ ⊂ ℝ^*p*^ denote the gene-expression spaces for *p* shared genes of the source and target domains𝒟^s^, respectively. In the source domain ^*s*^, samples are labeled as sensitive (*y*^*s*^ = 1) or resistant (*y*^*s*^ = 0),

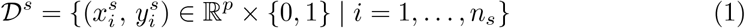

while in the target domain

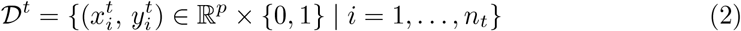

the labels 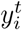 are unavailable or available only for a very small subset of the dataset. The distributions of the two domains are, in general, different.

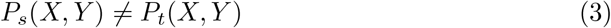

due to biological and technical differences between bulk and single-cell transcriptomes.

### 4.2 Domain adaptation methods

We benchmarked four representative architectures: SCAD [22], scDEAL [23], SSDA4Drug [26], and scATD (sf-VAE-dist) [24], alongside single-domain CatBoost models, serving as a baseline. Each domain-adaptation method aligns the source and target domains through distinct mechanisms, as explained below. An overview of the architectural paradigms, alignment mechanisms, and training regimes of all methods is provided in Fig. 6.

**Fig. 6:**
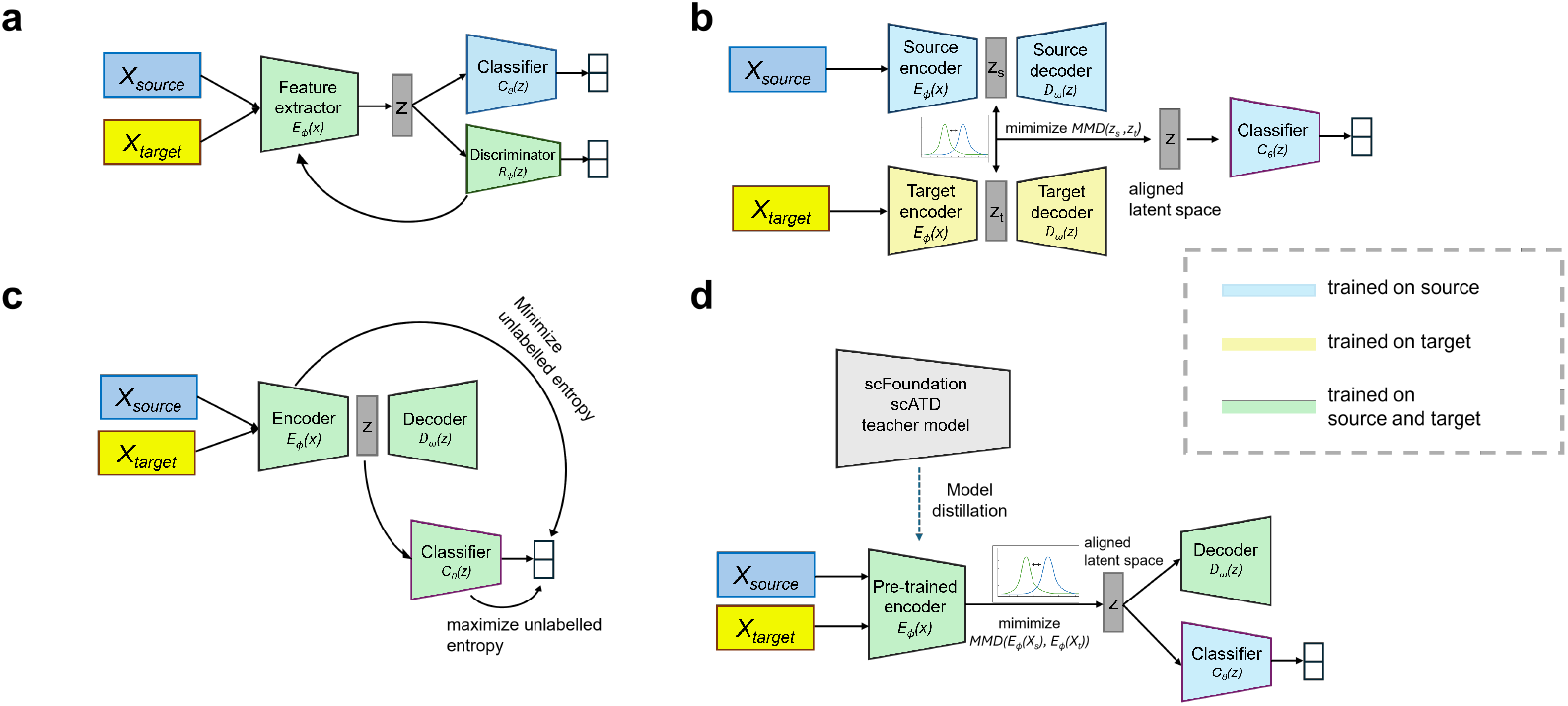
Schematic overview of the four benchmarked architectures. Blue and yellow blocks denote source (bulk) and target (single-cell) inputs, respectively; gray blocks indicate latent representations *z*. (a) **SCAD:** A shared encoder maps both domains into a common latent space, a classifier is trained on source labels, and an adversarial discriminator encourages domain-invariant features. (b) **scDEAL:** Separate source/target denoising autoencoders reconstruct inputs and an MMD loss aligns the two latent distributions while a shared classifier predicts drug response.(c) **SSDA4Drug:** Semi-supervised adaptation combining reconstruction, adversarial perturbations, and minimax entropy (MME) on unlabeled targets; min *H* and max *H* denote entropy minimization and maximization steps implemented via gradient reversal. (d) **scATD:** A scFoundation-trained teacher distills representations into a lightweight Res-VAE; the distilled encoder is used as a pretrained backbone and fine-tuned with an MMD loss to align source and target latents for downstream classification.

#### 4.2.1 SCAD – Adversarial domain alignment

SCAD [22] follows an adversarial domain adaptation paradigm based on adversarial discriminative domain adaptation (ADDA) [19] (see Fig. 6a). An encoder *E*_*ϕ*_, parameterized by *ϕ*, which is shared between source and target, maps input expression profiles to a common latent space *z* = *E*_*ϕ*_(*x*). A classifier *C*_*θ*_ (with parameters *θ*) classifies source domain features using latent representations, while a domain discriminator *R*_*ψ*_ (with parameters *ψ*) distinguishes the latent representations of source and target embeddings.

SCAD minimizes a supervised source classification loss

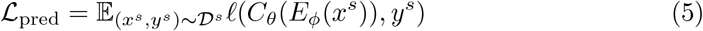

where ℓ(·,·) denotes the binary cross-entropy loss − [*y* log(ŷ) + (1 − *y*) log(1 − ŷ)], and simultaneously aligns latent spaces: while the discriminator learns to distinguish domains, the encoder learns to fool the discriminator by maximizing it, thereby removing domain-specific cues:

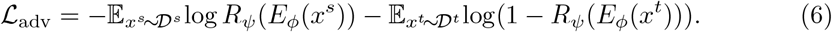

The overall loss function is

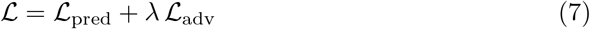

with a tunable adversarial loss weight *λ*. The adversarial game aims to generate domain-invariant features that allow the source trained predictor to generalize to the target domain.

#### 4.2.2 scDEAL – MMD-based alignment with dual autoencoders

In contrast to SCAD, which employs a single autoencoder to align domains, scDEAL [23] uses two separate denoising autoencoders (DAEs) for bulk and single-cell domains, coupled with a shared predictor. Each DAE reconstructs input expression profiles via its encoder–decoder pair *E*_*ϕ*_(·), *D*_*ω*_(·),

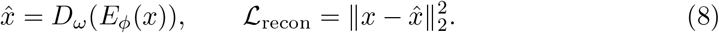

After independent DAE pretraining, domain adaptation is achieved by aligning the latent representations of the two encoders through a Maximum Mean Discrepancy (MMD) loss

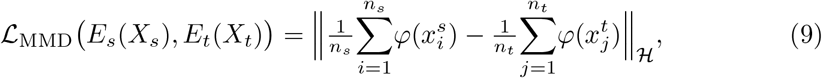

where φ(·) maps the data into a Reproducing Kernel Hilbert Space defined by the kernel *k*(*x, x*^*′*^) = ⟨φ(*x*), φ(*x*^*′*^)⟩_ℋ_. The overall objective jointly optimizes the encoders *E*_*s*_, *E*_*t*_ and the shared predictor *C*,

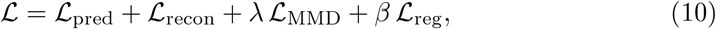

where ℒ_pred_ is the cross-entropy loss on bulk labels (Eq. 5) and ℒ_reg_ encourages local consistency among single-cell embeddings by maximizing cosine similarity within Louvain-defined cell clusters, thereby preserving intrinsic cell-type structure. Similarly to *λ* for ℒ_MMD_, the weight of the regularization loss ℒ_reg_ can be adjusted by tuning *β*. By minimizing Eq. (9), scDEAL learns domain-invariant representations that allow the bulk-trained predictor to generalize drug-response prediction to single-cell data (see Fig. 6b).

#### 4.2.3 SSDA4Drug – Semi-supervised minimax entropy adaptation

In contrast to the unsupervised methods above, SSDA4Drug [26] incorporates a small subset of labeled target samples to guide adaptation while leveraging unlabeled target data through a minimax conditional-entropy objective. Its encoder–decoder backbone follows the DAE notation introduced in Eq. (8), ensuring consistent feature extraction across domains. The shared encoder *E* and classifier *C* (with the final linear layer *W*_*P*_ serving as class prototypes) are trained with four complementary loss terms,

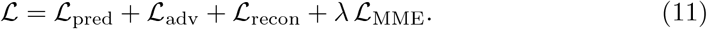

where *λ* is a hyperparameter used to balance the minimax entropy loss and the classification loss on the labeled samples. The reconstruction term ℒ_recon_ is the DAE loss defined in Eq. (8), reused here for consistency. The prediction lossℒ _pred_ extends Eq. (5) to include both source and few-shot target labels:

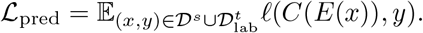

The adversarial robustness term ℒ_adv_ introduces subtle perturbations within the vicinity of the input data using gradient-based perturbations with KL divergence. Here, *p*(*y x*) denotes the class-posterior predicted by the classifier *C*(*E*(*x*)), and *q*(*y*) denotes the empirical label distribution, instantiated as a binary vector for labeled samples.

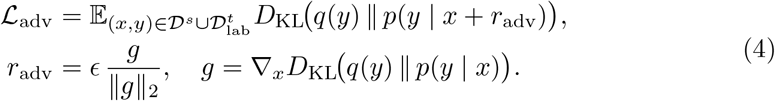

The main driver of domain adaptation in SSDA4Drug is the minimax conditional entropy (MME) objective applied to unlabeled target samples. For the unlabeled target set 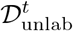, we define the conditional entropy

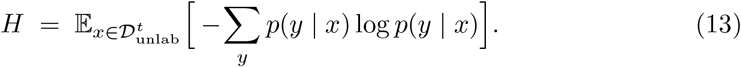

MME updates *E* and *C* adversarially:

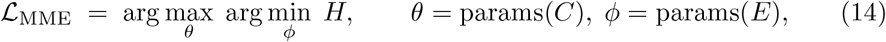

implemented with a gradient reversal layer.

Intuitively, the classifier first maximizes entropy on unlabeled targets, which pulls the class prototypes toward the centers of the target feature cloud (to prevent overfitting to the source boundary). Next, the feature extractor minimizes the same entropy, clustering target features around the prototypes and sharpening decision boundaries in the target domain. The alternation between these two objectives yields a discriminative yet domain-invariant feature representation under sparse target supervision [29].

#### 4.2.4 scATD – Foundation-model-guided MMD alignment

scATD [24] leverages pretrained single-cell foundation models to enable efficient, interpretable drug-response prediction. The variant used here, scATD-sf-dist, is based on the scFoundation model [25], which was first fine-tuned on Panglao single-cell data in the original study (scATD-sf) and subsequently distilled into a lighter student model (scATD-sf-dist). This distillation embeds the representational knowledge of scFoundation into a smaller residual variational autoencoder (Res-VAE) backbone through cosine-distance (COD) loss, preserving accuracy while reducing computational cost [24]. In our setup, we adopt the distilled Res-VAE encoder *E*_*ϕ*_ as a pretrained backbone and follow a two-stage training strategy. First, only the classifier head *C*_*θ*_ is trained on our labeled source data while keeping *E*_*ϕ*_ frozen. Next, the encoder is unfrozen and jointly fine-tuned with the classifier using the MMD alignment loss, continuing until the validation loss decreases, allowing for modest adaptation without overfitting.

The overall loss during fine-tuning combines prediction, reconstruction, and kernel-based alignment losses:

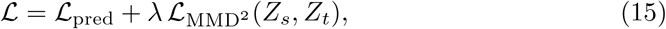

where ℒ_pred_ is the supervised cross-entropy on labeled bulk data (Eq. 5) and 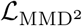 aligns bulk and single-cell latent features. Unlike scDEAL, where MMD compares RKHS-embedded features from two denoising encoders, scATD applies a squared MMD on the latent representations 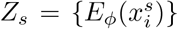 and 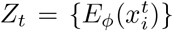 of the Res-VAE using a Gaussian kernel with a learnable scale, emphasizing direct statistical alignment in latent space:

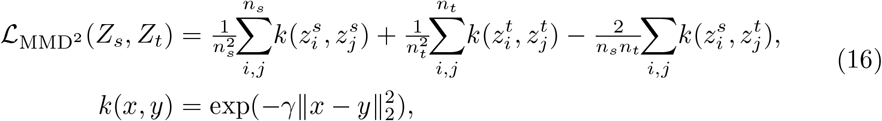

where *γ* is a learnable bandwidth parameter of the Gaussian kernel. This approach exploits foundation-model knowledge distilled into the Res-VAE while maintaining computational efficiency and stable cross-domain alignment.

#### 4.2.5 CatBoost – Non-adaptive gradient boosting baselines

To establish strong non-adaptive reference points, we trained gradient-boosted decision tree (GBDT) models using the CatBoost library [27]. CatBoost builds an ensemble of *T* oblivious decision trees by minimizing a log-loss objective through ordered boosting, thereby reducing target leakage and overfitting on small datasets. Given input features *x* and binary labels *y*∈{0, 1}, the prediction function is

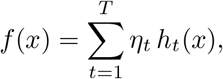

where each weak learner *h*_*t*_ is a binary decision tree, and *η*_*t*_ is its learning rate. The optimization minimizes the empirical risk

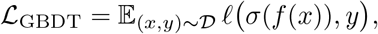

with ℓ the binary cross-entropy loss and *σ*(·) the logistic sigmoid. Two CatBoost variants were evaluated: (i) Source-only: trained exclusively on labeled bulk data ^*s*^, representing a non-adaptive baseline under covariate shift; (ii) Few-shot: trained on 𝒟^*s*^ augmented with three labeled single-cells from the target domain per class from 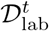, providing a simple few-shot comparison. Since CatBoost does not explicitly align source and target distributions, these models quantify the performance achievable from supervised learning alone under domain shift.

### 4.3 Code

All implementations, as published in their GitHub repositories, were refactored into a unified PyTorch Lightning (v2.5.5) training stack with consistent logging, early stopping, and metric computation to ensure full reproducibility, available through our GitHub repository (https://github.com/cbg-ethz/SC-Bulk-Domain-Adaptation). To ensure consistency with the original code, we tested our Lightning implementation under fixed conditions. We found that it either matches or slightly exceeds the performance of the original implementations (Supplementary Fig. 10).

### 4.4 Data sources

#### 4.4.1 Source domain data (cell lines, bulk transcriptomes)

The source domain consisted of cell line expression data from the Genomics of Drug Sensitivity in Cancer (GDSC) [9]. For 625 GDSC cell lines, RNA-seq data was available from Garcia *et al*. 2018 [30] and Klijn *et al*. 2015 [31], for the remaining 332 GDSC cell lines without publicly available RNA-seq data, we used the microarray expression data of the GDSC cancer cell line panel (release 8.3, June 2020). We downloaded the pre-processed RMA-normalized basal expression profiles (Affymetrix Human Genome U219 array) from https://www.cancerrxgene.org/.

The RNA-Seq count matrix of Garcia *et al*. 2018 was obtained from the European Genome-Phenome Archive (EGAD00001001357) and the count matrix of Klijn *et al*. 2015 [31] was obtained from the EMBL-EBI Expression Atlas (E-MTAB-2706).

As a drug response measure, we employ the binarized *C*_max_ viability obtained from [12]. The *C*_max_ viability denotes the cell line viability rates at *C*_max_, which is the peak plasma concentration of each drug after the administration of the highest clinically recommended dose. Labels were assigned as *sensitivity* = 1 − *C*_max_ viability and used as binarized labels, setting the threshold at *C*_max_ ≥0.5, so that sensitive cells are labeled with 1 and resistant cells with 0. Since label binarization helped in some cases but hurt performance in others, we defined a tunable Boolean hyperparameter for binarization.

#### 4.4.2 Target domain data (single-cell transcriptomes)

We downloaded and processed 19 single-cell RNA-Seq datasets for which cells were treated with 10 different drugs. This includes datasets used in the original publications of SCAD, scDEAL, SSDA4Drug, and scATD if drug-specific *C*_max_ viability labels for the source data were available. The number of benchmark datasets is greater than in the publications of scDEAL (6), SCAD (10), SSDA4Drug (10), and scATD (16).

Most publicly available single-cell RNA-Seq drug-response datasets, including those used in prior benchmarking studies assign response labels based on treatment status. In particular, untreated cells are simply labeled as sensitive, while cells surviving drug exposure are labeled as resistant. This approach ignores the presence of pre-existing resistant subclones in the untreated population and conflates intrinsic resistance with transcriptional changes caused by treatment. As a result, treatment status rather than drug susceptibility often dominates the signal, leading to an increased separation of “sensitive” and “resistant” cells in the expression space (Fig. 3b,c). In addition, the RNA-seq data from the SCC47 and JHU006 cell lines in GSE157220, used in the SCAD and SSDA4Drug publications, were labeled according to the epithelial senescence-related (EpiSen) program [32]. The EpiSen score captures transcriptional programs associated with cellular senescence and has been shown to correlate with drug-sensitivity and resistance across multiple compounds. In the original study, Kinker *et al*. [32] isolated EpiSen-high and EpiSen-low cells by fluorescence-activated cell sorting (FACS) based on the expression of the marker genes Claudin4 and AXL. Cells with EpiSen scores within the top 10% of all cells were labeled as EpiSen-high, and those within the bottom 10% as EpiSen-low. Although these labels are not just a proxy of the treatment status, the resulting sensitivity–resistance separation is artificially amplified because only the most extreme 20% of cells were labeled, leaving the remaining 80% unlabeled and thus excluded from the analysis.

To counteract such biases, we also incorporated datasets in which resistance was inferred independently of treatment survival. Specifically, we included pre-treatment datasets where resistance was assigned through lineage tracing, such as GSE223003 (Olaparib) and GSE111014 (Afatinib) (Fig. 3a). Unlike post-treatment labeling, lineage tracing labels cells with DNA barcodes to track their clonal relationships over time, linking pre-treatment transcriptional states to eventual survival or resistance outcomes. This allows resistance labels to be assigned to cells prior to drug exposure, providing a biologically more faithful reference for evaluating cross-domain generalization.

We also tested the models on two xenograft datasets with drug-sensitive and resistant post-treatment samples: GSE175716 and GSE228154, and a dataset with labeled pre-treatment samples from chronic lymphocytic leukemia patients. Table 1 provides an overview of all 19 datasets and their respective dataset IDs for access in the Gene Expression Omnibus (GEO) or the Broad Institute’s single-cell portal.

**Table 1:**
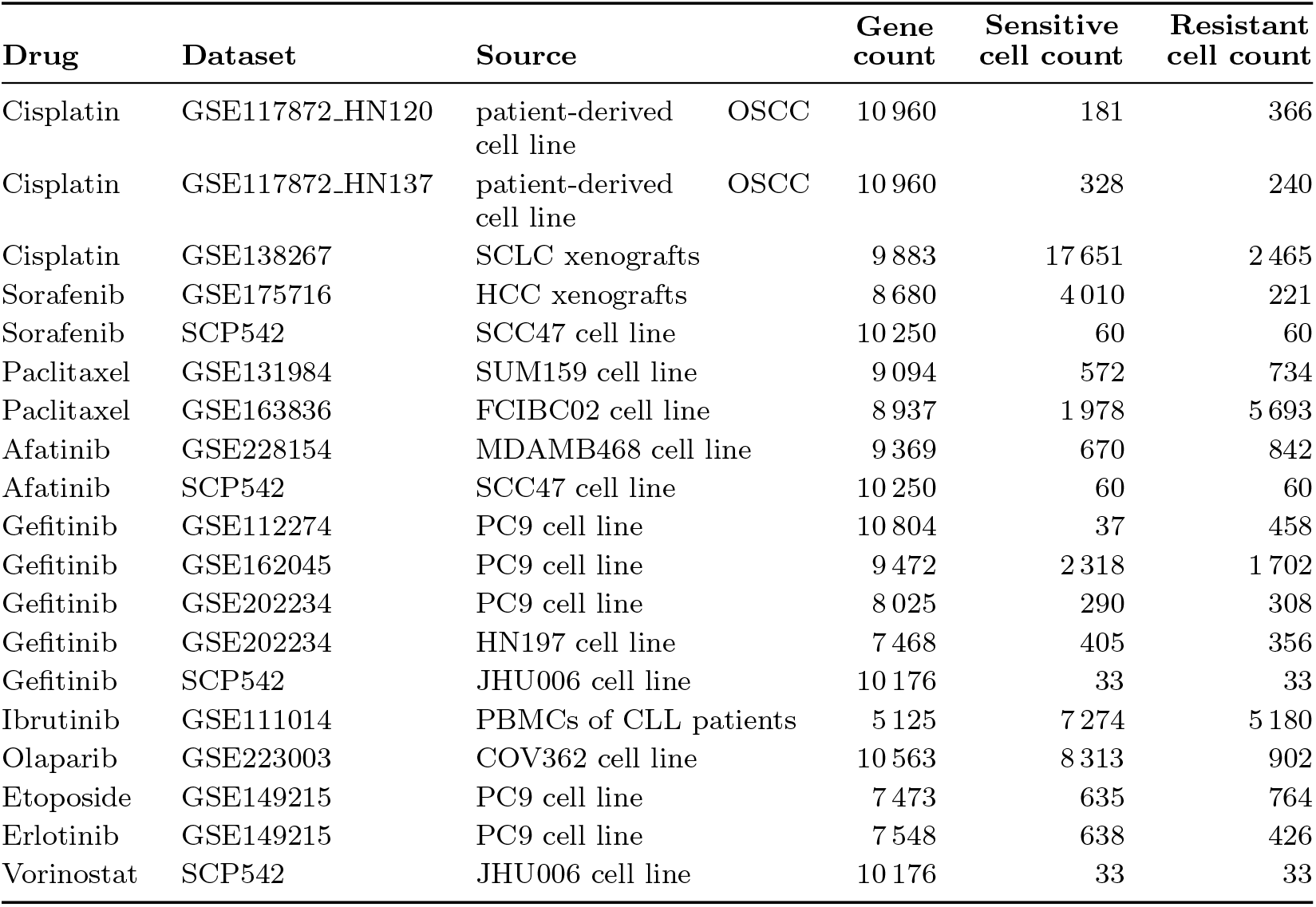
Target domain datasets.

### 4.5 Data processing

GDSC expression data was downloaded in an RMA-normalized state and not further processed. Raw RNA-Seq counts from Garcia *et al*. 2018 [30] were normalized to counts per million (CPM) and subsequently log_10_(CPM + 1)-transformed. Singlecell RNA-sequencing datasets were processed using Scanpy (v1.11.4). First, cells expressing fewer than 500 genes, containing more than 10% mitochondrial transcripts, or exhibiting more than 40% ribosomal content were removed. In addition, genes detected in fewer than 3% of all cells were removed. Furthermore, outlier cells, with median expression levels greater than the dataset median plus three standard deviations, were discarded. Lastly, all datasets were normalized to counts per million and log_10_(CPM + 1)-transformed.

#### 4.5.1 Gene-set alignment

For all models, bulk and single-cell feature spaces were aligned through the intersection of gene symbols across datasets, with the number of shared genes ranging from 5,125 to 12,316. For scATD sf-dist, which requires a fixed vocabulary of 19,264-genes, missing genes were zero-imputed for both source and target, as in the original publication [24].

### 4.6 Benchmark framework and evaluation

#### 4.6.1 Splits and class balance

For source domain data, a split of 64% for training, 16% for validation, and 20% for testing, stratified by the labels, was applied. Target domain data were split into stratified training (80%) and testing (20%) subsets. SSDA4Drug additionally required a few (3 per class by default) labeled target samples. These data points were drawn exclusively from the target-train partition. To address class imbalance, weighted over-sampling of the minority class was employed for every model. In contrast to some of the original implementations, synthetic data generation techniques such as SMOTE or other augmentation strategies were not used.

#### 4.6.2 Thresholding and Metrics

For each trained model and dataset, the decision threshold is chosen to maximize MCC on the source–validation set. We evaluated all models on held–out source–test and target–test partitions using the following metrics: MCC, AUROC, AUPRC, accuracy, precision, recall, and specificity. The complete set of metric values, including F1-Score, accuracy, specificity, sensitivity, and MCC, is summarized in Supplementary Figure 7.

#### 4.6.3 Hyperparameter Optimization

In contrast to the original fixed-epoch method implementations, all domain-alignment stages were trained for up to 100 epochs with patience = 10, monitored on the source-validation loss. For multi-stage methods (scDEAL, scATD), the initial domain-specific stages ran for 50 epochs. Learning-rate schedulers, dropout, and optimizers follow each method’s original defaults unless overridden by hyperparameter search.

We perform per-dataset Bayesian optimization with Optuna (100 trials per model–dataset pair; see Supplementary Methods for parameter ranges and search protocol). The objective was source validation-set MCC after per-trial threshold tuning. The best trial is selected based on the MCC score; if multiple trials are tied, the highest AUROC is used, followed by the AUPRC. In cases of trials with identical best-source validation MCC, AUROC, and AUPRC, the mean target performance metrics across all tied runs for the source and target test sets were reported.

### 4.7 Independent target evaluation

To assess the generalization ability of each method beyond the datasets seen during training, we conducted independent target evaluations. For each drug, models were trained on the source and primary target domains using hyperparameters optimized for the source domain MCC, and were subsequently evaluated on one or more independent single-cell datasets measuring resistance or sensitivity to the same compound.

Before training, we intersected the gene sets of the source, target, and independent target datasets to ensure that only features shared across all domains were used. For four drugs, two independent target datasets were available: for Cisplatin, three (6 distinct target–independent target evaluation pairs), and for Gefitinib, five (20 distinct target–independent target evaluation pairs). Each independent target dataset was used exclusively for testing without further partitioning into training or validation subsets. Detailed descriptions of hyperparameter optimization, threshold calibration, and reproducibility settings are provided in the Supplementary Methods (Section 5.2).

## Declarations

### Ethics approval and consent to participate

Not applicable

### Consent for publication

All authors have read and approved the final manuscript and consent to its upload (preprinting) on bioRxiv.

### Data availability

All datasets used in this study are publicly available. Bulk cell line expression and drug-response data were obtained from the Genomics of Drug Sensitivity in Cancer (GDSC, release 8.3, December 2020; https://www.cancerrxgene.org) and from the RNA-seq datasets of Garcia *et al*. [30] (EGA accession EGAD00001001357) and Klijn *et al*. [31] (https://www.ebi.ac.uk/gxa/experiments/E-MTAB-2706/Results). scRNAseq datasets were downloaded from the NCBI Gene Expression Omnibus (GEO) under the accession numbers listed in Table 1. scRNA-Seq data for the JHU006 and SCC47 cell lines was downloaded from the single-cell portal (https://singlecell.broadinstitute.org/singlecell/study/SCP542/).

The preprocessed benchmark datasets, together with all training configurations and results, are available on Zenodo under https://zenodo.org/records/17868777.

No proprietary or patient-identifiable data were used in this study.

### Code availability

Our code is available at https://github.com/cbg-ethz/SC-Bulk-Domain-Adaptation.

### Author contributions

K.L. conceived the study. K.L. and M.E-M. designed and supervised the study. M.B. implemented the methods and conducted the experiments. M.E-M. and M.B. provided the initial draft of the manuscript. M.B., M.E.-M. and K.L. wrote and edited the manuscript. N.B. reviewed and edited the manuscript. All authors analyzed and interpreted the data and results.

## Acknowledgement

We thank Lea Eckhart for carefully proofreading our manuscript and testing our code base.

## 5 Supplementary

### 5.1 Supplementary Results

**Fig. 7:**
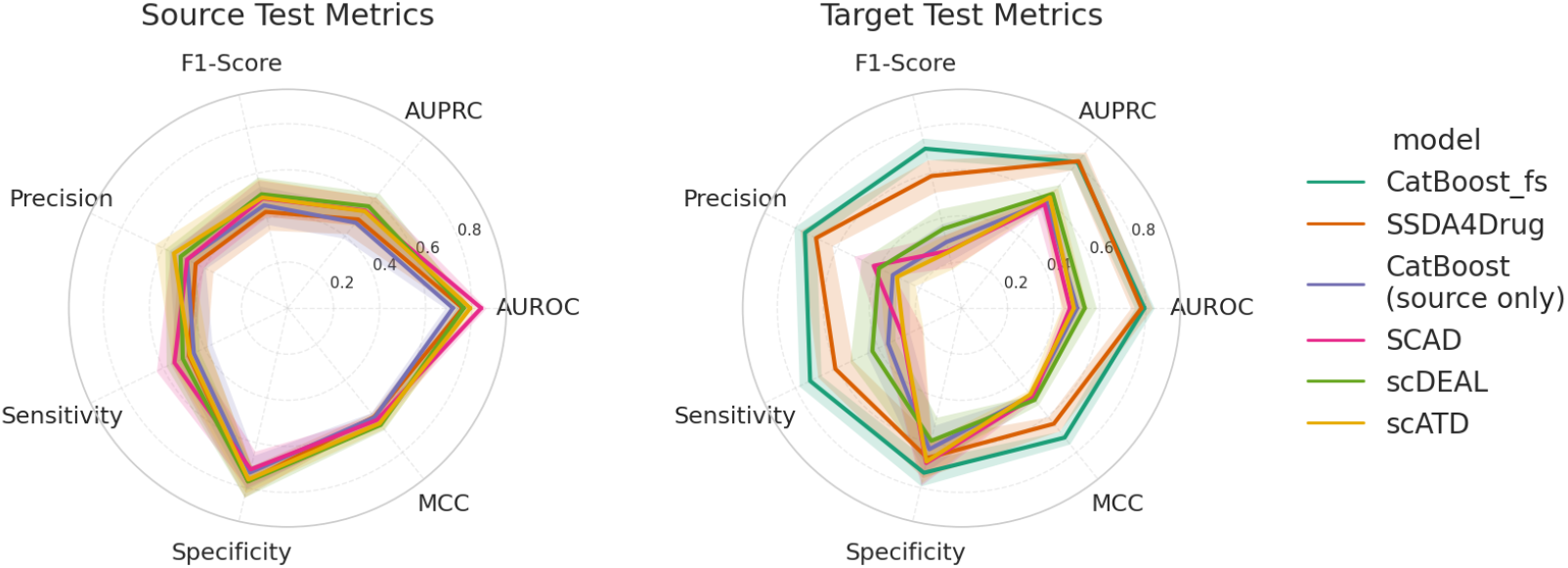
Complete evaluation of all domain adaptation methods and baseline models. Lines indicate the mean and opaque areas around the mean line indicate the standard error of the mean.

**Fig. 8:**
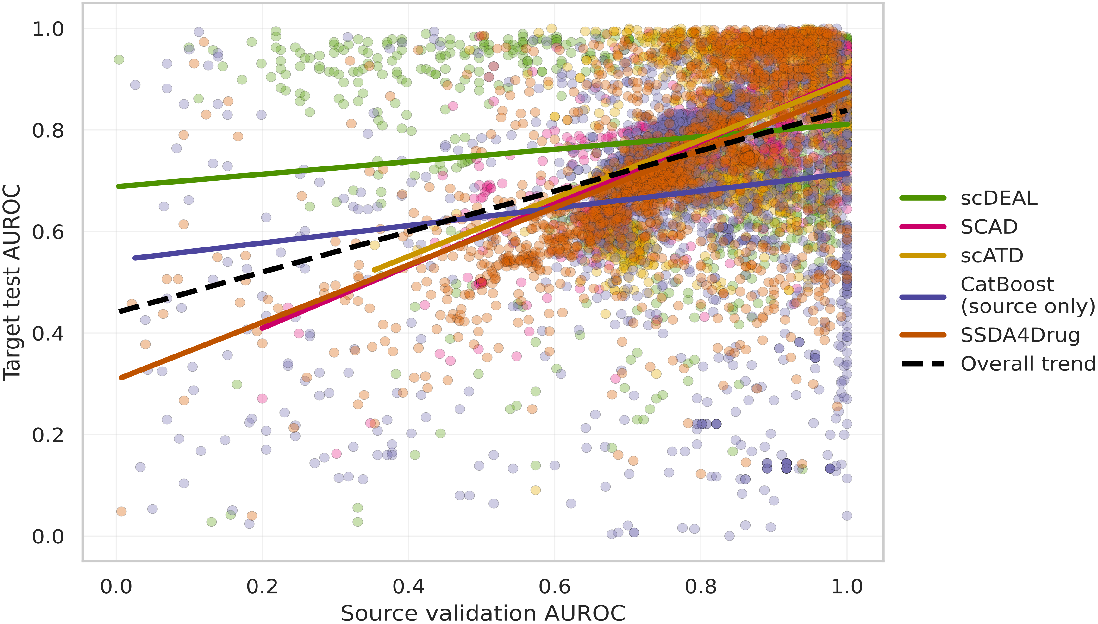
Association between source validation and source test AUROC. Points represent trained models and trendlines depict linear regression estimates.

**Fig. 9:**
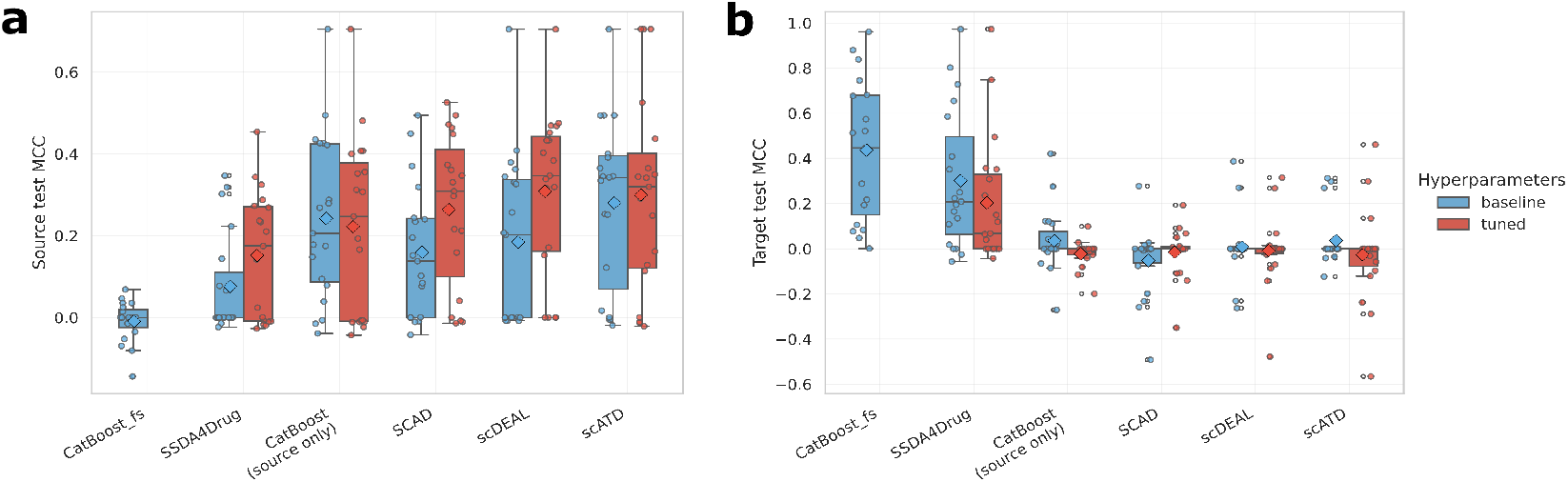
Baseline and tuned MCC scores. (a) Source MCC, (b) target MCC. Diamonds depict means.

### 5.2 Supplementary Methods

### 5.3 Data processing

We used bulk RNA-seq data from cancer cell lines as the source domain and drug-treated single-cell RNA-seq (scRNA-seq) data as the target domain.

#### 5.3.1 Data harmonization and cleaning

Gene expression matrices were loaded for all datasets, and gene identifiers were standardized to Ensembl IDs during preprocessing. Genes with zero expression across all samples were removed, and duplicate gene entries were collapsed by retaining the one with the highest mean expression. The resulting gene sets were intersected across domains to define a unified feature space for model training.

#### 5.3.2 Dataset partitioning

Source-domain data were split into training (64%), validation (16%), and test (20%) subsets, stratified by drug sensitivity to preserve class balance. Target-domain data were partitioned into training (80%) and test (20%) splits, stratified by resistance labels. The source-validation split was used exclusively for early stopping, hyperparameter optimization, and threshold selection to prevent target-data leakage.

### 5.3.3 Feature scaling

For all models requiring normalized input, we fitted a StandardScaler (Scikit-learn) on the source-training partition only. The same scaling transformation was then applied to all remaining partitions (source validation, source test, target train, and target test). This ensured consistent input scaling while avoiding any information leakage from target or test sets.

### 5.4 Validation of Lightning re-implementation

To verify that the PyTorch Lightning refactor preserved model behavior, we bench-marked our implementations against the original repositories using identical data splits, seeds and either default (scDEAL, scATD, SSDA4Drug) or author-specified (SCAD) optimal hyperparameters. For these tests, we also did not use our own processed data but the processed data provided by the SCAD authors (which is identical to the data provided by the SSDA4Drug authors)

We did not apply any custom sampling schemes, such as undersampling or SMOTE but instead applied weighted, oversampling of the minority class for all models.

Across all evaluated datasets, the Lightning versions achieved performance comparable to, or slightly better than, the original implementations, confirming faithful reproduction of the training logic. On average, the Lightning implementation performed slightly better (mean AUROC: 0.7141, median AUROC: 0.7594, mean AUPRC: 0.7671, median AUPRC: 0.7993) than the original implementations (mean AUROC: 0.7073, median AUROC 0.6813, mean AUPRC: 0.7604, median AUPRC: 0.7694)

### 5.5 Hyperparameter tuning

To identify optimal hyperparameters for each domain-adaptation framework, we conducted systematic searches using the Optuna optimization library. For each drug–dataset pair, we performed an independent Optuna study per model, running 100 trials. We employed the Tree-structured Parzen Estimator (TPE) sampler, chosen for its efficiency in exploring high-dimensional search spaces.

The optimization objective was to maximize the Matthews Correlation Coefficient (MCC) on the source validation set. After each training run, the classification threshold was determined dynamically by scanning probability cutoffs between 0.05 and 0.95 and selecting the one maximizing the Matthews correlation coefficient (MCC) on the source-validation set. All threshold-sensitive reported metrics, inclucing accuracy, precision, recall, specificity, sensitivity, and MCC were computed using this optimized threshold.

**Fig. 10:**
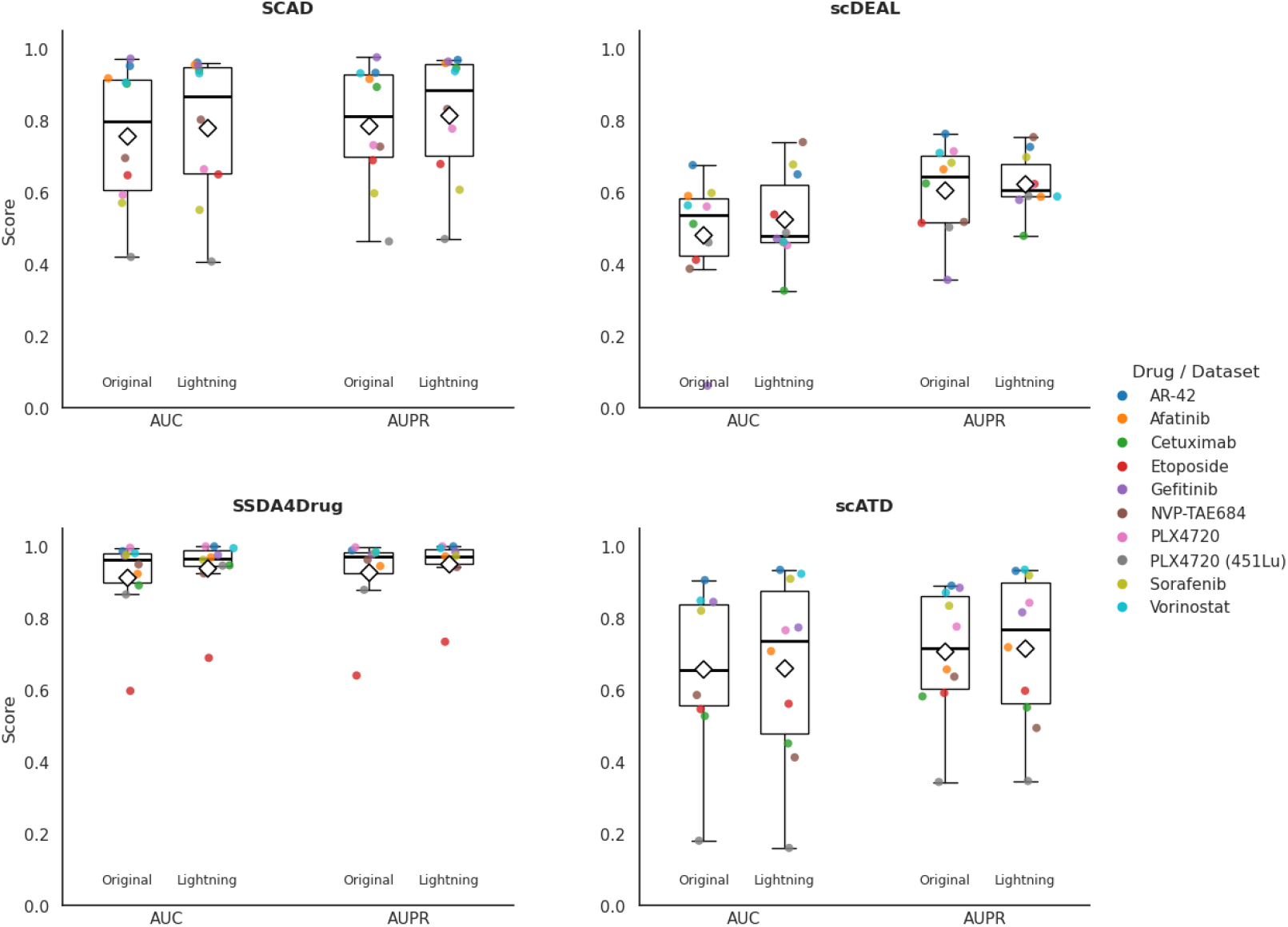
Head-to-head comparison of the original SCAD (a), scDEAL (b), SSDA4Drug (c), and scATD (d) implementations versus our PyTorch Lightning refactor. The performance is indicated by AUROC and AUPRC scores and the data, splits, and hyperparameters were used as documented in the SCAD authors’ GitHub repository. For scATD, scDEAL, and SSDA4Drug, default hyperparameters were used.

All runs were tracked using the Weights & Biases (W&B) platform to ensure full reproducibility. Logged artifacts included hyperparameter configurations, training/-validation curves, and final evaluation metrics on the source validation, source test, and target test sets. The complete experiment logs are available in Source Data XYZ. To ensure deterministic behavior across frameworks, we fixed a global random seed of 42 for all relevant libraries (`NumPy, PyTorch, Optuna`), controlling for random initialization, data shuffling, and split assignment.

The search grids were defined to be at least as comprehensive as those used in the original publications or their released source code. For frameworks with incomplete documentation (**SSDA4Drug, scATD**), parameter ranges were inferred based on common conventions and model architecture. Baseline hyperparameters (shown in bold in Table 2) correspond to either the default values in the authors’ repositories (**SSDA4Drug, scDEAL**) or, when multiple were reported, the most frequently used settings in the supplementary materials. In cases of missing or contradictory documentation (e.g., discrepancies between GitHub README files and Supplementary Tables), we estimated reasonable defaults based on comparable configurations in related models.

**Table 2:** Unified hyperparameter search grids grouped by model.

| Model | Hyperparameter | Candidate values | Baseline value |
| --- | --- | --- | --- |
| **SCAD** | lr | floats in [1e-4, 5e-2] | 5e-4 |
|  | dropout | floats in [0.1, 0.5] | 0.5 |
|  | h_dim | integers in [512, 1024] (step 128) | 1024 |
|  | predictor_z_dim | integers in [128, 256] (step 64) | 128 |
|  | mbS | {8, 16, 32, 64} | 8 |
|  | mbT | {8, 16, 32, 64} | 8 |
|  | lam1 | floats in [0.1, 10.0] (log-uniform) | 1 |
| **SSDA4Drug** | lr | floats in [5e-5, 1e-2] (log-uniform) | 1e-3 |
|  | dropout | floats in [0.1, 0.5] | 0.3 |
|  | encoder_h_dims | {"256,128", "512,256", "1024,512"} | "512,256" |
|  | predictor_h_dims | {"64,32", "128,64", "256,128"} | "64,32" |
|  | batch_size | {64, 128, 256} | 128 |
|  | epsilon | 0 or floats in [1e-4, 1e-2] (log-uniform) | 0 |
|  | lambda | floats in [0.05, 0.2] | 0.1 |
| **scDEAL** | lr | floats in [1e-3, 1e-1] (log-uniform) | 1e-2 |
|  | dropout | floats in [0.1, 0.5] | 0.3 |
|  | encoder_h_dims | {"256,256", "512,256", "512,512"} | "512,256" |
|  | predictor_h_dims | {"16,8", "32,16", "64,32", "128,64"} | "64,32" |
|  | bottleneck | integers in [64, 512] (step 64) | 128 |
|  | mmd_weight | floats in [0.1, 1.0] | 0.25 |
| **scATD** | lr | floats in [1e-5, 5e-2] (log-uniform) | 1e-4 |
|  | mmd_weight | floats in [0.1, 1.0] | 1 |
|  | batch_size | {32, 64, 128, 256} | 128 |
|  | weight_decay | floats in [1e-6, 1e-2] (log-uniform) | 1e-3 |

### 5.6 Runtime analysis

Table 3 summarizes the average training time per model, measured across all drugs and datasets. Neural network-based domain-adaptation models exhibited considerable variance in execution time due to multi-stage training and adversarial losses, while gradient-boosting baselines ran efficiently.

**Table 3:** Training time on a Nvidia Tesla V100 GPU with tuned hyperparameter configurations. Mean, median, and standard deviation across 21 datasets.

| Model | mean (s) | median (s) | SD (s) |
| --- | --- | --- | --- |
| CatBoost source | 18.7 | 17.0 | 9.9 |
| CatBoost few shot | 0.9 | 1.0 | 0.6 |
| scDEAL | 400.0 | 155.5 | 717.1 |
| SCAD | 55.1 | 40.5 | 54.5 |
| SSDA4Drug | 30.2 | 11.0 | 61.2 |
| scATD | 92.9 | 93.5 | 87.0 |

**Table 4:** Independent target datasets used for cross-dataset evaluation.

| Dataset (GEO) | Drug | Data type / cell line |
| --- | --- | --- |
| GSE117872 (HN120) | Cisplatin | OSCC cell line (patient-derived) |
| GSE117872 (HN137) | Cisplatin | OSCC cell line (patient-derived) |
| GSE138474 | Cisplatin | SCLC xenografts (8 PDXs) |
| GSE175716 | Sorafenib | HCC xenograft |
| GSE157220 (SCC47) | Sorafenib | HNSCC cell line |
| GSE131984 (SUM159) | Paclitaxel | TNBC cell line |
| GSE163836 (FCIBC02) | Paclitaxel | Breast cancer cell line |
| GSE228154 (MDAMB468) | Afatinib | TNBC cell line |
| GSE157220 (SCC47) | Afatinib | HNSCC cell line |
| GSE112274 (PC9) | Gefitinib | Lung cancer cell line |
| GSE162045 (PC9) | Gefitinib | Lung cancer cell line |
| GSE202234 (PC9/H1975/JHU006) | Gefitinib | Lung/HNSCC cell lines |

### 5.7 Independet Target datasets

The datasets below were used for our target/independent-target transfer experiments.

### 5.8 Statistical independence of dataset features and performance

**Table 5:** Statistical test of metadata-performance associations.

| Association | Test | Estimate (p-value) |
| --- | --- | --- |
| drug – AUROC (fs models) | Kruskal–Wallis H-test | $H = 4.27$ ($p = 0.893$) |
| shared genes – AUROC (all models) | Spearman | $\rho = 0.148$ ($p = 0.180$) |
| shared genes – AUROC (fs models) | Spearman | $\rho = -0.104$ ($p = 0.597$) |
| sensitivity fraction – AUROC (all models) | Spearman | $\rho = -0.056$ ($p = 0.612$) |
| sensitivity fraction – AUROC (fs models) | Spearman | $\rho = 0.071$ ($p = 0.718$) |

**Table 6:** Datasets and evaluation metrics of the tested domain adaptation frameworks. **Color legend:** Black = used exclusively in one study or only in this benchmark. blue = used in SCAD, SSDA4Drug, and this benchmark. orange = used in scATD and this benchmark. purple = used in scDEAL, scATD, and this benchmark. red = used in SCAD, SSDA4Drug, and scATD (not in this benchmark).

| Model | Published evaluation metrics | Datasets used for benchmarking |
| --- | --- | --- |
| **SCAD** [22] | AUROC, AUPRC | **10 in total:** **GSE149215**, **SCP542_JHU006** (labels for 3 drugs), **SCP542_SCC47** (labels for 4 drugs), **GSE108383_451Lu**, **GSE108383_A375**. |
| **scDEAL** [23] | AUROC, AUPRC, F1-Score | **6 in total:** **GSE117872_HN120**, **GSE117872_HN137**, **GSE112274**, **GSE149383**, **GSE110894**. |
| **SSDA4Drug** [26] | AUROC | **10 in total:** **GSE149215**, **SCP542_JHU006** (labels for 3 drugs), **SCP542_SCC47** (labels for 4 drugs), **GSE108383_451Lu**, **GSE108383_A375**. |
| **scATD** (sf-VAE-dist) [24] | **Main benchmark:** AUROC, AUPRC, F1-Score. **Independent target benchmark:** AUROC, AUPRC, F1-Score, Precision, Recall, Accuracy, MCC | **16 in total:** **GSE131984**, **GSE163836**, **GSE202234**, **GSE117872_HN120**, **GSE117872_HN137**, **GSE112274**, **GSE108383_A375**, **GSE108383_451Lu**, **GSE223779**, **GSE149214**, **GSE149383**, **GSE186960**, **GSE111014**, **GSE169246**, **GSE108397**. |
| **Our benchmark** | AUROC, AUPRC, Accuracy, Precision, Recall, F1-Score, MCC | **21 in total:** **GSE149215_Etop**, **GSE149215_Erlo**, **SCP542_JHU006** (labels for 2 drugs), **SCP542_SCC47** (labels for 2 drugs), **GSE131984**, **GSE163836**, **GSE202234_PC9**, **GSE202234_HN1975**, **GSE111014**, **GSE117872_HN120**, **GSE117872_HN137**, **GSE112274**, **GSE140440_PC3**, **GSE138267**, **GSE175716**, **GSE228154**, **GSE162045**, **GSE223003**. |

